# MutCleaner: Cleaning and Standardizing Biological Mutation Datasets for Variant Effect Prediction

**DOI:** 10.64898/2026.09.06.749687

**Authors:** Ziyu Shi, Yuxiang Tang, Mengxin Yang, Shize Yu, Yancheng Shi, Yunxin Xu

**Author notes:** These authors contributed equally (Ziyu Shi, Yuxiang Tang).

## Abstract

**Summary:** Protein mutation datasets are widely used in variant effect prediction and protein engineering, but datasets from different sources often lack consistent conventions for mutation representation, sequence representation, data organization, and experimental labels, making these resources difficult to integrate directly and limiting their use in downstream analyses and modeling. MutCleaner is an extensible Python framework that cleans, validates, and standardizes protein- and codon-level mutation datasets through composable cleaning pipelines, unified sequence and mutation data structures, and dataset-specific cleaners. It provides standardized resources covering 16 protein and codon mutation datasets with more than 12.14 million mutation records

**Availability and implementation:** MutCleaner is an open-source Python package released under the Apache License 2.0. The source code is available on GitHub, the package is distributed through PyPI, and the documentation is available online.

## 1 Introduction

Amino acid mutations can affect protein stability, molecular interactions, and function, thereby altering organismal phenotypes or contributing to disease processes [1, 2]. Quantitative measurement of protein mutation effects is therefore fundamental to understanding protein sequence–function relationships. Protein mutation effects can be measured individually for single or a limited number of variants using biochemical or biophysical approaches, such as chemical denaturation and enzyme kinetics assays [3], or quantified in parallel across large variant libraries using multiplexed assays of variant effect (MAVE). Among these high-throughput experimental strategies, deep mutational scanning (DMS) and massively parallel reporter assay (MPRA) are representative approaches [4]. The mutation-effect data generated by these experiments can be used for training and evaluating predictive models, as well as for downstream applications such as protein engineering, disease mechanism studies, and drug design.

However, differences across studies in experimental design, data-processing workflows, and data-recording practices result in substantial heterogeneity among existing protein mutation datasets, including differences in data structures, field names, and annotation schemes. Raw datasets such as the mega-scale protein folding stability dataset [5], the TrpB combinatorial mutation fitness dataset [6], the human myoglobin mutation stability dataset [7], the CTX-M *β*-lactamase mutation-effect dataset [8], the Human Domainome site-saturation mutagenesis dataset [9], and higher-order combinatorial mutation datasets for protein stability [10], among others [11–19], may use different file formats and field names and represent mutation annotations, sequence information, and experimental measurements in different ways. These datasets may also contain missing values, duplicate records, invalid mutation annotations, and inconsistencies between mutation annotations and sequence information. Together, these differences and data-quality issues hinder direct dataset integration and downstream analysis.

Preprocessing of protein mutation data currently relies largely on scripts written for individual datasets. Although this approach offers flexibility, cleaning steps and error information are often not organized or recorded in a standardized manner, making such workflows difficult to reuse, inspect, and extend. Public resources such as MaveDB [20] and ProteinGym [21] have collected, curated, and released large amounts of mutation-effect data. MaveDB focuses primarily on the storage, sharing, and management of multiplexed variant effect measurements, whereas ProteinGym provides standardized protein mutation-effect datasets and benchmark resources for evaluating variant effect prediction models. Although these resources have substantially improved the accessibility and usability of existing data, different resources still lack consistent conventions for aspects such as field naming and mutation representation. Meanwhile, new experimental datasets continue to be generated, and their raw formats often do not directly conform to the data representations used by existing resources, requiring researchers to develop dataset-specific cleaning scripts. A general cleaning and standardization framework is therefore still needed that encapsulates common data-cleaning operations as reusable steps and allows these steps to be configured and combined for different datasets. A detailed comparison of ProteinGym, MaveDB, and MutCleaner in terms of their primary purposes, mutation annotation standardization, data validation, preprocessing extensibility, and structured outputs is provided in Supplementary Table S5. To address this need, we developed MutCleaner, an extensible Python framework for reproducible cleaning, validation, and standardization of protein and codon mutation datasets from heterogeneous sources and formats. Through composable cleaning pipelines, MutCleaner validates mutation annotations and sequences, checks mutation–sequence consistency during data cleaning, and separately exports valid and failed records. We further applied MutCleaner to 16 protein and codon mutation data resources and provide standardized resources containing more than 12.14 million mutation records.

## 2 Design and implementation

For new protein or codon mutation datasets, MutCleaner uses a modular design to assemble reusable cleaning steps and dataset-specific cleaning steps into ordered cleaning pipelines. These pipelines clean and validate raw datasets and ultimately export standardized datasets with a consistent structure.

### 2.1 Unified Sequence and Mutation Representations

To validate mutation annotations and sequences, MutCleaner defines unified classes for DNA, RNA, and protein sequences, as well as for amino acid and codon mutations. MutCleaner supports mutation-effect data at both the protein level and the DNA/RNA codon level and establishes a unified representation and validation framework around reference sequences, mutation annotations, and mutated sequences (Fig 1A). DNA, RNA, and protein sequence strings are instantiated as sequence objects of the corresponding types, and their validity is checked during object construction according to the alphabet constraints defined for each sequence type. Raw mutation strings are normalized during instantiation with respect to position indexing and delimiters for multiple mutations, while mutation symbols, positions, and mutation combinations are also validated. On this basis, MutCleaner performs successive validation of data types, sequences, mutation annotations, and mutation–sequence consistency, as illustrated by the core validation workflow in Fig 1B. Beyond normalization and validity checking, MutCleaner also supports sequence generation, inference of mutation annotations, conversion of codon mutations to amino acid mutations, and translation of DNA or RNA sequences. Different datasets can select the appropriate operations according to the information available and their specific cleaning requirements. For example, when any two of the reference sequence, mutation annotation, and mutated sequence are available, the third can be derived using the corresponding operations, and consistency among the three can subsequently be validated. When mutation annotations are provided at the codon level but an amino-acid-level representation is required, the corresponding representation conversion can also be performed.

**Figure 1.**
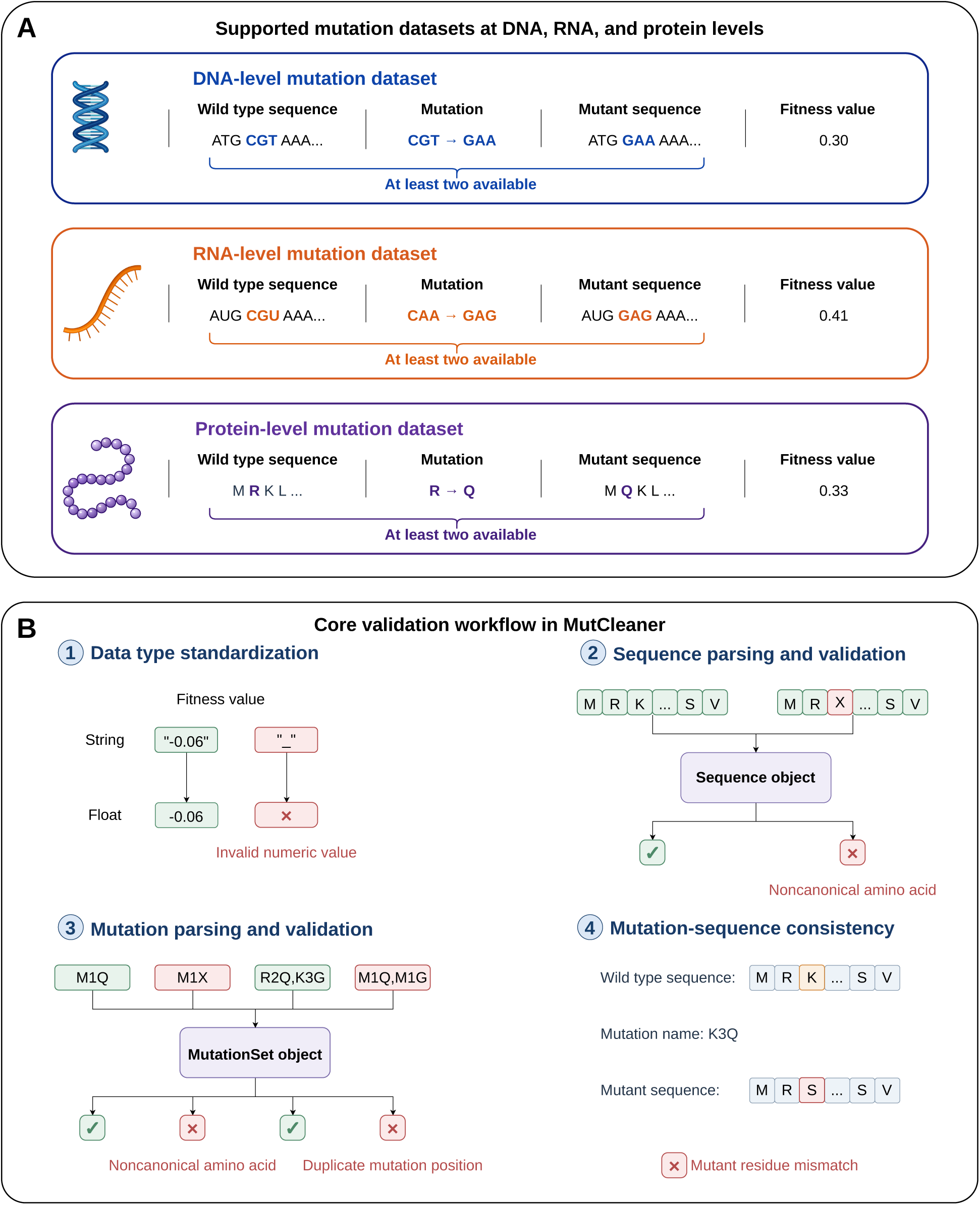
Overview of mutation data representation and validation workflow in MutCleaner. **A,** Supported mutation-effect datasets at the DNA, RNA, and protein levels. **B,** Core validation steps for data types, sequences, mutation annotations, and mutation–sequence consistency.

### 2.2 Composable Cleaning Pipeline and Standardized Outputs

MutCleaner decomposes the data-cleaning process into a series of processing steps and combines them in an explicit order to form dataset-specific cleaning pipelines. Users can configure reusable processing steps according to dataset characteristics and incorporate additional dataset-specific steps when necessary. Dataset-specific workflows can also be defined through configuration classes such as BaseCleanerConfig. These configuration classes specify dataset-dependent parameters while reusing common cleaning operations, thereby reducing the repetitive development effort required to integrate new datasets. For processing steps that may generate invalid records, records that pass validation or are processed successfully continue through the main data flow, whereas records that fail validation or processing are retained as artifacts for subsequent data tracking and error analysis. In practical dataset curation, MutCleaner can identify and record multiple classes of data-quality issues, including invalid mutation annotations, invalid sequences, mutation–sequence inconsistencies, non-numeric labels, and missing values in required fields. Dataset-level quality-control statistics are provided in Supplementary Table S2. The pipeline also records step names, execution status, runtime, and exception information to support inspection of pipeline execution and error diagnosis. For computationally intensive operations, including mutation annotation validation and inference and mutated-sequence generation, performance benchmarks show that increasing the number of workers can further reduce overall runtime; detailed runtime changes and speedup values are provided in Supplementary Fig S2 and Supplementary Table S3.

Cleaned datasets are encapsulated as MutationDataset objects, providing a consistent representation for mutation data from different sources together with built-in validation and dataset summary functions. For example, before downstream analysis, MutationDataset can validate mutation positions and the consistency between mutation annotations and reference sequences. During export, MutCleaner creates a separate directory for each reference sequence according to its reference_id. Each directory contains data.csv, which stores mutation annotations, mutated sequences, and labels; wt.fasta, which stores the wild-type sequence; and metadata.json, which records reference-sequence metadata and summary statistics. This standardized organization facilitates direct use of the processed data in downstream variant effect prediction tasks (Supplementary Fig S1).

## 3 Dataset Coverage and Standardized Resources

The data resources currently curated with MutCleaner consist primarily of protein mutation datasets and also include codon mutation datasets, covering protein stability, fitness and epistasis, protein interactions, and codon mutation effects. The resource currently comprises 16 datasets containing more than 12.14 million mutation records in total and can support mutation-effect analysis and model evaluation across datasets of different scales (Supplementary Table S1). The datasets were obtained from published studies, public data resources, and user-provided datasets. The input files required by the cleaning pipelines are hosted on Hugging Face. MutCleaner supports an end-to-end workflow from raw-data download, cleaning, and standardization to output storage, enabling reproducibility and traceability of the resulting standardized datasets. The generated standardized datasets can be used directly as consistent inputs for downstream mutation-effect analyses. To further evaluate the reliability of the data-cleaning and standardization results, we compared MutCleaner outputs for the cDNA-display proteolysis dataset with the corresponding datasets independently curated by ProteinGym. Across 64 matched DMS assays, the mutation-effect values showed high agreement between the two resources, with mean Spearman and Pearson correlation coefficients of approximately 0.9996 and a mean absolute label difference of 0.0032 (Supplementary Table S4).

## 4 Conclusion

In summary, MutCleaner provides a modular and reusable framework for transforming heterogeneous raw mutation data into validated and consistently structured standardized resources. The project also provides a large collection of curated and validated standardized datasets that can be directly used for downstream mutation-effect analyses. Although datasets from different sources still differ in data structures, field naming, and annotation schemes and therefore require additional dataset-specific processing steps, MutCleaner provides a framework and reusable tools for continuously expanding standardized mutation data resources and facilitating downstream mutation-effect analyses.

## Author contributions

Y.X. conceived the study, proposed the methodology and developed the overall research framework. Y.X. and Y.T. designed the initial framework, implemented the initial codebase, and conducted preliminary testing. Z.S. optimized the implementation details and integrated the final datasets.

Y.T. curated the Human Domainome Dataset, ProteinGym DMS Substitutions Dataset, Protein cDNA Proteolysis Dataset, ddG Dataset and dTm Dataset. Z.S. curated the ArchStabMS1E10 Epistasis Dataset, Antitoxin ParD3 Epistasis Dataset, TrpB Epistasis Dataset, Protein Human Myoglobin Epistasis Dataset, Codon Human Myoglobin Epistasis Dataset and CTXM Epistasis Dataset. M.Y. curated the MGnify ddG Dataset. Y.S. curated the RBD ACE2 Dataset and RBD Antibody Dataset. S.Y. curated the Codon cDNA Proteolysis Dataset and Codon DMS Substitutions Dataset.

Y.X. and Z.S. analyzed the results and wrote the manuscript. All authors reviewed and approved the final manuscript.

## Data availability

The mutation datasets used in this study are publicly available from the MutCleaner dataset repository on Hugging Face. These datasets, together with the cleaning workflows provided by MutCleaner, enable reproduction of the standardized datasets described in this study.

## S1 Software architecture

MutCleaner is organized into three main modules: mutcleaner.cleaners, mutcleaner.core, and mutcleaner.utils. The cleaners module defines reusable data-cleaning steps and dataset-specific cleaning pipelines; the core module implements MutCleaner’s core biological data structures and pipeline framework; and the utils module provides supporting utilities for data conversion, parallel processing, raw-data retrieval, and input/output operations.

The main software structure of MutCleaner is as follows:

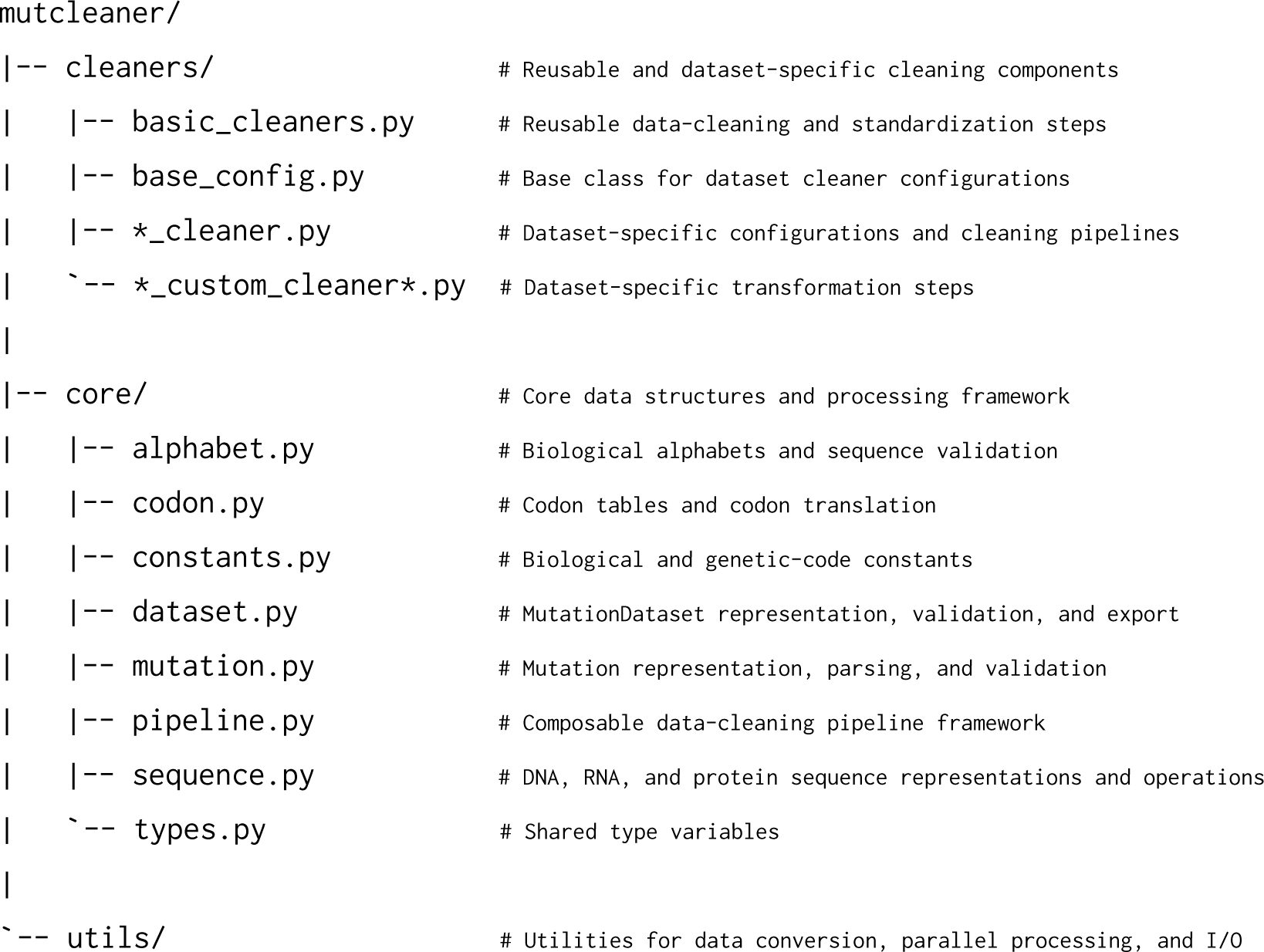

### S1.1 mutcleaner.cleaners

The cleaners module implements both reusable data-cleaning steps and dataset-specific cleaning workflows.

basic_cleaners.py contains general data-processing operations that can be reused across datasets, including dataset loading, column operations, data filtering and cleaning, data type conversion, mutation annotation validation, sequence and mutation processing, label processing, and conversion to the standardized MutCleaner mutation dataset format.

base_config.py defines the abstract configuration base class used by dataset-specific cleaners and provides common configuration parameters, configuration validation, serialization and deserialization to and from dictionary and JSON formats, and configuration merging.

The *_cleaner.py module for each dataset defines the dataset-specific configuration parameters and complete cleaning pipeline.

These modules combine reusable cleaning steps from basic_cleaners.py with dataset-specific transformation steps in a predefined order to construct a complete cleaning workflow for the corresponding raw dataset.

Processing operations that are specific to an individual dataset and have limited reuse across datasets are defined separately in the corresponding *_custom_cleaner*.py modules.

### S1.2 mutcleaner.core

The core module contains the core biological data structures and data-processing framework used by MutCleaner.

alphabet.py defines the biological alphabets used for DNA, RNA, and protein sequences and provides functions for checking sequence characters and validating sequence validity.

codon.py implements codon-table representations and translation from codons to amino acids, together with functions for identifying start and stop codons.

constants.py stores biological constants shared across MutCleaner, including standard and ambiguous nucleotide symbols, standard and ambiguous amino acid symbols, base-complement relationships, mappings between one-letter and three-letter amino acid codes, and the standard genetic code tables for DNA and RNA.

dataset.py defines the MutationDataset data structure, which provides a unified representation of the relationships among mutation sets, reference sequences, and experimental labels. This module also provides dataset validation, calculation of summary statistics, data representation conversion, and export to the standardized MutCleaner dataset format.

mutation.py defines representations for amino acid substitution mutations and codon substitution mutations, together with their corresponding mutation sets, and implements mutation annotation parsing, validation, and mutation type classification.

pipeline.py implements the composable data-cleaning pipeline framework used to manage the execution order of cleaning steps, immediate and delayed execution, step execution status and runtime, and auxiliary outputs generated during data cleaning.

sequence.py defines data abstractions for DNA, RNA, and protein sequences and provides sequence validation, mutation application, and conversion and processing operations associated with different sequence types.

types.py defines shared type variables used throughout MutCleaner to provide consistent type constraints for sequences, mutations, mutation sets, cleaner configurations, and related objects.

### S1.3 mutcleaner.utils

The utils module provides supporting functionality required by the cleaning pipelines and core data structures, including parallel execution of cleaning tasks, construction of standardized datasets, label parsing, mutation annotation and data type conversion, sequence input/output, data-source information management, and downloading of raw data files.

## S2 Dataset background and cleaning procedures

The datasets included in MutCleaner originate from diverse experimental systems and research objectives and differ substantially in experimental design, mutation types, phenotypic measurements, and data organization. To facilitate understanding of the biological context of each dataset and its associated cleaning workflow, this section first briefly describes the data source, experimental measurements, and label definitions for each dataset, and then summarizes the main cleaning and standardization steps performed by MutCleaner.

### S2.1 Human Domainome Dataset

Beltran et al. constructed the Human Domainome 1 Supplementary Table 2 Dataset by performing large-scale site-saturation mutagenesis across hundreds of human protein domains and systematically measuring the effects of amino acid substitutions on intracellular protein abundance using an abundance protein-fragment complementation assay (aPCA) [1]. Supplementary Table 2 summarizes the filtered single-amino-acid substitutions and their normalized aPCA fitness values. This metric reflects the effects of amino acid mutations on intracellular protein abundance and protein folding stability. The main cleaning and standardization procedures for the Human Domainome Supplementary Table 2 Dataset were as follows:

1. Use the normalized_fitness field as the mutation-effect label;
2. Remove records containing stop mutations or missing values in key fields;
3. Subtract the sequence offset encoded in the domain_ID field from each mutation position and combine the wild-type and mutant amino acids to construct the mutation annotation;
4. Validate the constructed mutation annotations;
5. Infer the wild-type sequence from the mutation annotation and mutated sequence, and verify the consistency of the inferred wild-type sequences within the same domain;
6. Convert the cleaned data into the standardized MutCleaner format.

Beltran et al. further developed a thermodynamic model using aPCA measurements from multiple homologous domains within the same protein family in Human Domainome 1 to infer mutation-associated changes in folding free energy (ΔΔ*G*) that are relatively conserved across family members [1]. Supplementary Table 4 therefore contains model-inferred stability measurements rather than results from an independent mutational screening experiment. The main cleaning and standardization procedures for the Human Domainome Supplementary Table 4 Dataset were as follows:

1. Use the mean_kcalmol_scaled field as the mutation-effect label;
2. Subtract the domain start position coordinates encoded in the PFAM_entry field from each mutation position to obtain the relative position within the domain, and combine the wild-type and mutant amino acids to construct the mutation annotation;
3. Add the corresponding full-length wild-type sequence according to uniprot_ID, and extract the corresponding domain sequence using the domain start and end positions;
4. Apply the mutation annotation to the domain wild-type sequence to generate the corresponding mutated sequence;
5. Convert the cleaned data into the standardized MutCleaner format.

### S2.2 ProteinGym DMS Substitutions Dataset

Notin et al. developed ProteinGym, a large-scale DMS benchmark for evaluating protein variant effect prediction and protein design methods, integrating mutation-effect data from diverse proteins and experimental systems [2]. The DMS_score standardizes the direction of mutation-effect measurements across assays such that higher scores correspond to higher variant fitness, while the specific biological interpretation of the score depends on the original experiment. The main cleaning and standardization procedures for the ProteinGym DMS Substitutions Dataset were as follows:

1. Use the DMS_score field as the mutation-effect label;
2. Validate and standardize mutation annotations;
3. Infer the wild-type sequence from the mutation annotation and mutated sequence, and verify the consistency of wild-type sequences within the same protein;
4. Convert the cleaned data into the standardized MutCleaner format.

### S2.3 Protein cDNA Proteolysis Dataset

Tsuboyama et al. developed cDNA-display proteolysis technology, in which proteins are covalently linked to their encoding cDNA and subjected to proteolysis at different protease concentrations, followed by deep sequencing to enable high-throughput inference of the thermodynamic folding stability of small protein domains [3]. The resulting resource contains systematic single-mutant scans of natural and designed proteins as well as a subset of combinatorial mutations. The dG_ML field represents quality-filtered estimates of folding stability Δ*G*, whereas ddG_ML represents quality-filtered estimates of mutation-associated stability changes ΔΔ*G*. The main cleaning and standardization procedures for the Protein cDNA Proteolysis Dataset were as follows:

1. For the Protein cDNA Proteolysis dG Dataset, use the dG_ML field as the mutation-effect label; for the Protein cDNA Proteolysis ddG Dataset, use the ddG_ML field as the mutation-effect label;
2. Convert the ddG_ML field from string to numeric type and remove records that cannot be converted;
3. Use the mut_type field as the mutation annotation, and validate and standardize the mutation annotations;
4. Average the labels of duplicate records with the same protein and mutation annotation;
5. Infer the wild-type sequence from the mutation annotation and mutated sequence, and verify the consistency of wild-type sequences within the same protein;
6. Convert the cleaned data into the standardized MutCleaner format.

### S2.4 ddG Dataset

The ddG Dataset was curated by Xu et al. for training and evaluating GeoDDG and comprises the S8754, S669, S461, S783, and M1261 protein stability datasets [4]. ΔΔ*G* describes mutation-associated changes in protein folding free energy, and the datasets additionally retain experimental conditions such as pH and temperature. The main cleaning and standardization procedures for the ddG Dataset were as follows:

1. Use the ddG field as the mutation-effect label;
2. Infer mutation annotations by comparing the wild-type and mutated sequences;
3. Group replicate measurements with the same protein and mutation annotation and preferentially select the record measured at the experimental pH closest to 7.0. If multiple records have the same distance from pH 7.0, select the record measured at the temperature closest to 25 ^◦^C as the final label;
4. Convert the cleaned data into the standardized MutCleaner format.

### S2.5 dTm Dataset

The dTm Dataset was curated by Xu et al. for training and evaluating GeoDTm and contains protein thermostability data integrated from existing protein thermostability databases [4]. Δ*T*_m_ describes the mutation-induced change in protein melting temperature, and the dataset also retains experimental conditions such as pH. The main cleaning and standardization procedures for the dTm Dataset were as follows:

1. Use the dTm field as the mutation-effect label;
2. Infer mutation annotations by comparing the wild-type and mutated sequences;
3. Group replicate measurements with the same protein and mutation annotation and select the record measured at the experimental pH closest to 7.0 as the final label;
4. Convert the cleaned data into the standardized MutCleaner format.

### S2.6 ArchStabMS1E10 Epistasis Dataset

Faure et al. constructed multiple combinatorial mutation libraries for GRB2-SH3 and SRC and measured the effects of different mutation combinations on intracellular protein abundance and ligand binding using AbundancePCA and BindingPCA [5]. Supplementary Table 4 summarizes variant fitness values estimated from the sequencing data using DiMSum, with fitness values from different experiments representing the corresponding experimental phenotypes, including protein abundance and binding. The main cleaning and standardization procedures for the ArchStabMS1E10 Epistasis Supplementary Table 4 Dataset were as follows:

1. Use the fitness field as the mutation-effect label;
2. Infer mutation annotations by comparing the wild-type and mutated sequences for each protein;
3. Convert the cleaned data into the standardized MutCleaner format.

Based on the fitness measurements obtained from the combinatorial mutagenesis experiments described above, Faure et al. further used the MoCHI thermodynamic model to infer mutation-associated changes in protein folding and binding free energies, together with energetic coupling between pairs of mutations [5]. Supplementary Table 5 therefore summarizes model-inferred changes in folding and binding free energy and energetic coupling rather than directly measured experimental fitness. The main cleaning and standardization procedures for the ArchStabMS1E10 Epistasis Supplementary Table 5 Dataset were as follows:

1. Retain records with conf=True;
2. Add the corresponding wild-type sequence according to the library identifier;
3. Use the mean_kcal/mol field as the mutation-effect label;
4. Validate mutation annotations;
5. Correct mutation positions affected by the positional offset in library 4 by adding 3 to the corresponding mutation positions;
6. Add the original label of each double-mutant record to the labels of its corresponding single mutants to obtain the final label for the double mutant, and remove double-mutant records for which the corresponding single-mutant records are unavailable;
7. Apply the mutation annotations to the wild-type sequence to generate the mutated sequences;
8. Convert the cleaned data into the standardized MutCleaner format.

### S2.7 Antitoxin ParD3 Epistasis Dataset

This dataset originates from combinatorial mutagenesis studies of the ParD–ParE toxin–antitoxin system. Lite et al. constructed a complete library containing all 20^3^ = 8,000 amino acid combinations at three key interaction positions in the ParD3 antitoxin and quantified the toxin-neutralization capacity of different variants using cellular growth competition and deep sequencing [6]. Ding et al. subsequently used this library for further combinatorial mutagenesis studies and data analysis [7]. The version used by MutCleaner was obtained from the processed variant-effect data subsequently released in the CoVES study [8]. The main cleaning and standardization procedures for the Antitoxin ParD3 Epistasis Dataset were as follows:

1. Use the label field as the mutation-effect label;
2. Add the wild-type sequence according to the original publication;
3. Validate mutation annotations;
4. Average replicate measurements of the same mutation and subtract the wild-type label to obtain the relative mutation effect;
5. Apply the mutation annotations to the wild-type sequence to generate mutated sequences;
6. Convert the cleaned data into the standardized MutCleaner format.

### S2.8 TrpB Epistasis Dataset

Johnston et al. constructed a complete 20^4^ = 160,000 combinatorial mutation landscape across four residues in the active site of a thermostable TrpB and measured variant fitness using a pooled-culture enrichment assay in tryptophan-auxotrophic *E. coli* [9]. The fitness values therefore quantify TrpB catalytic function through cellular growth. The main cleaning and standardization procedures for the TrpB Epistasis Dataset were as follows:

1. Use the fitness field as the mutation-effect label;
2. Supplement the wild-type sequence according to the original study;
3. Validate and standardize mutation annotations;
4. Average repeated measurements for identical mutations and subtract the wild-type label to obtain the relative mutation effect;
5. Apply mutation annotations to the wild-type sequence to generate mutated sequences;
6. Convert the cleaned data into the standardized MutCleaner dataset format.

### S2.9 Protein Human Myoglobin Epistasis Dataset

Küng et al. used yeast surface display combined with fluorescence-activated cell sorting (FACS) and deep sequencing to measure the expression levels of human myoglobin variants at large scale, yielding an expression fitness score [10]. This metric is strongly correlated with the thermostability of soluble human myoglobin, and the dataset contains single mutations, double mutations, and a small number of higher-order combinatorial mutations. The main cleaning and standardization procedures for the Protein Human Myoglobin Epistasis Dataset were as follows:

1. Use the fitness field as the mutation-effect label;
2. Add the wild-type sequence according to the original publication;
3. Convert codon-level mutations into amino-acid-level mutations and remove records involving stop codons;
4. Validate and standardize amino acid mutation annotations and convert mutation positions to 0-based indexing;
5. Apply the mutation annotations to the wild-type sequence to generate mutated sequences;
6. Average replicate measurements of the same mutation;
7. Convert the cleaned data into the standardized MutCleaner format.

### S2.10 CTXM Epistasis Dataset

Judge et al. systematically introduced pairwise combinatorial mutations across 17 active-site residues of the CTX-M-14 *β*-lactamase and measured variant fitness under cefotaxime and ampicillin selection using *E. coli* growth selection and deep sequencing [11]. The fitness values represent the relative fitness of each mutant compared with the wild type under the corresponding antibiotic selection condition, and the dataset contains large numbers of single- and double-mutant records. The main cleaning and standardization procedures for the CTXM Epistasis Dataset were as follows:

1. Retain records with is.reads0=True and expand each original pairwise record into two single-mutant records and one double-mutant record, retaining the corresponding fitness label for each;
2. Add the wild-type sequence according to the original publication;
3. Map mutation positions from the Ambler numbering system to the corresponding positions in the actual protein sequence;
4. Validate and standardize mutation annotations and convert mutation positions to 0-based indexing;
5. Average duplicate records with the same mutation annotation;
6. Apply the mutation annotations to the wild-type sequence to generate mutated sequences;
7. Convert the cleaned data into the standardized MutCleaner format.

### S2.11 RBD ACE2 Dataset

The RBD ACE2 Dataset integrates multiple deep mutational scanning studies performed in different SARS-CoV-2 RBD variant backgrounds [12–15]. These studies used yeast-surface-display RBD mutant libraries together with concentration gradients of monomeric human ACE2, FACS sorting, and deep sequencing to obtain binding curves for individual variants and quantify their ACE2-binding affinity. Because the dataset covers multiple viral variant backgrounds, measurements across different RBD backgrounds can also be used to compare how mutation effects change during SARS-CoV-2 evolution. The main cleaning and standardization procedures for the RBD ACE2 Dataset were as follows:

1. Use the log10Ka field as the mutation-effect label;
2. Remove records with missing label values;
3. Validate and standardize mutation annotations, standardize the delimiter between multiple mutations to a comma, and convert mutation positions to 0-based indexing;
4. Average replicate measurements with the same RBD target and mutation annotation and subtract the corresponding wild-type label;
5. Add the corresponding wild-type sequence according to the RBD target;
6. Apply the mutation annotations to the wild-type sequence to generate mutated sequences;
7. Convert the cleaned data into the standardized MutCleaner format.

### S2.12 RBD Antibody Dataset

The RBD Antibody Dataset integrates multiple studies of antibody-binding escape from SARS-CoV-2 RBD mutations [16–18]. These studies used yeast-display RBD mutant libraries and FACS to enrich variants with reduced binding to serum antibodies or monoclonal antibodies, followed by deep sequencing to quantify antibody-binding escape associated with individual mutations. The analyses additionally incorporated pre-selection sequencing counts, RBD expression, and ACE2-binding measurements for quality filtering to reduce nonspecific escape signals caused by insufficient RBD expression or misfolding. The main cleaning and standardization procedures for the RBD Antibody Dataset were as follows:

1. Use the score field as the mutation-effect label;
2. Remove records that fail the pre-count and ACE2-binding/expression quality filters or have missing label values;
3. Validate and standardize mutation annotations, standardize the delimiter between multiple mutations to a comma, and convert mutation positions to 0-based indexing;
4. Average replicate measurements with the same RBD wild-type sequence, antibody, and mutation annotation and subtract the corresponding wild-type label;
5. Add the corresponding wild-type sequence according to the RBD reference and apply the mutation annotations to generate mutated sequences;
6. Convert the cleaned data into the standardized MutCleaner format.

### S2.13 MGnify ddG Dataset

Cho et al. used cDNA-display proteolysis to measure the absolute folding stability of 1.8 million diverse protein domains, primarily derived from the MGnify metagenomic database and ranging from 60 to 80 amino acids in length [19]. These domains span more than 200,000 sequence families, resulting in the large-scale MGnify Stability Dataset. Because the original dataset provides only the absolute folding stability Δ*G* of each sequence, an initial ΔΔ*G* dataset was generated for variant-effect prediction tasks. MMseqs2 was used to perform homology clustering with sequence-identity thresholds dynamically determined according to sequence length, while strictly ensuring that each cluster member differed from the cluster center, which was treated as the wild type (WT), by no more than 10 mutations. Clusters containing more than five mutated sequences were subsequently retained. The ΔΔ*G* values were then calculated from the difference between the Δ*G* of each mutated sequence and that of its corresponding WT. The main cleaning and standardization procedures for the MGnify ddG Dataset were as follows:

1. Use the ddG field as the mutation-effect label;
2. Infer mutation annotations by comparing the wild-type and mutated sequences;
3. Average duplicate records with the same protein and mutation annotation;
4. Convert the cleaned data into the standardized MutCleaner format.

### S2.14 Codon cDNA Proteolysis Dataset

The Codon cDNA Proteolysis Dataset was derived from the large-scale cDNA-display proteolysis resource established by Tsuboyama et al. [3] and retains mutation information at the DNA codon level together with the corresponding fitness measurements to describe the experimental effects associated with different codon mutations. The main cleaning and standardization procedures for the Codon cDNA Proteolysis Dataset were as follows:

1. Use the fitness field as the mutation-effect label;
2. Validate and standardize DNA codon mutation annotations;
3. Add the corresponding wild-type DNA sequence according to the protein name;
4. Average duplicate records with the same protein and codon mutation;
5. Apply the codon mutations to the wild-type DNA sequence to generate mutated sequences;
6. Convert the cleaned data into the standardized MutCleaner format.

### S2.15 Codon DMS Substitutions Dataset

The Codon DMS Substitutions Dataset is primarily derived from multiplexed assays of variant effect (MAVE) datasets deposited in MaveDB. MaveDB collects large-scale variant-effect measurements from diverse experimental systems and organizes variant information, experimental measurements, target sequences, and associated metadata as score sets [20]. To construct a resource suitable for codon-level mutation-effect analysis, protein-coding score sets from MaveDB were first screened and preprocessed before being incorporated into MutCleaner.

During preprocessing, only score sets with reliable coding sequences and without splice variants were retained. HGVS nucleotide variants were parsed against the corresponding wild-type coding sequences to derive codon substitutions and their associated amino acid substitutions, and the translated consequences were further checked against protein-level variant annotations. Records with reference-sequence mismatches, stop codons, insertions or deletions, or ambiguous coordinate systems were excluded. Experimental measurements were based on the scores provided by MaveDB and, when explicit wild-type records were available, were transformed into mutation effects relative to the wild type. Different score sets were further deduplicated according to relationships among their wild-type sequences, shared mutations, and correlations between their labels to reduce redundant inclusion of highly overlapping experimental data. The retained records were finally represented uniformly as DNA codon substitutions and used as input for the Codon DMS Substitutions Dataset. The main cleaning and standardization procedures for the Codon DMS Substitutions Dataset were as follows:

1. Read the mutation data and wild-type DNA sequence for each assay and use the assay name as the identifier for the corresponding reference sequence;
2. Use the label field as the mutation-effect label;
3. Validate and standardize DNA codon mutation annotations in the codon_mutation field;
4. Remove records in which the mutation introduces a stop codon;
5. Apply codon mutations to the corresponding wild-type DNA sequences to generate mutated sequences and validate consistency between the mutation annotations and reference sequences;
6. Convert the cleaned data into the standardized MutCleaner format.

### S2.16 Codon Human Myoglobin Epistasis Dataset

The Codon Human Myoglobin Epistasis Dataset and the Protein Human Myoglobin Epistasis Dataset are derived from the same human myoglobin deep mutational scanning experiment. Küng et al. used yeast surface display combined with FACS and deep sequencing to measure the expression levels of human myoglobin variants at large scale, yielding an expression fitness score that is strongly correlated with the thermostability of soluble human myoglobin [10]. Unlike the Protein Human Myoglobin Epistasis Dataset, in which the original codon mutations are converted into amino acid substitutions, the Codon Human Myoglobin Epistasis Dataset retains mutations at the codon level and uses the DNA sequence as the reference sequence, thereby providing a codon-level mutation-effect dataset. The main cleaning and standardization procedures for the Codon Human Myoglobin Epistasis Dataset were as follows:

1. Use the fitness field as the mutation-effect label;
2. Validate mutation annotations and convert mutation positions to 0-based indexing;
3. Remove records in which the mutation introduces a stop codon;
4. Apply codon mutations to the wild-type DNA sequence to generate mutated sequences and validate consistency between the mutation annotations and reference sequence;
5. Average replicate records corresponding to the same codon mutation;
6. Convert the cleaned data into the standardized MutCleaner format.

## S3 Standardized output format

MutCleaner organizes cleaned and standardized data using a consistent directory structure. For each reference sequence, a separate subdirectory is created according to its reference_id, containing three main output files: data.csv, wt.fasta, and metadata.json (Supplementary Fig S1).

**Figure S1.**
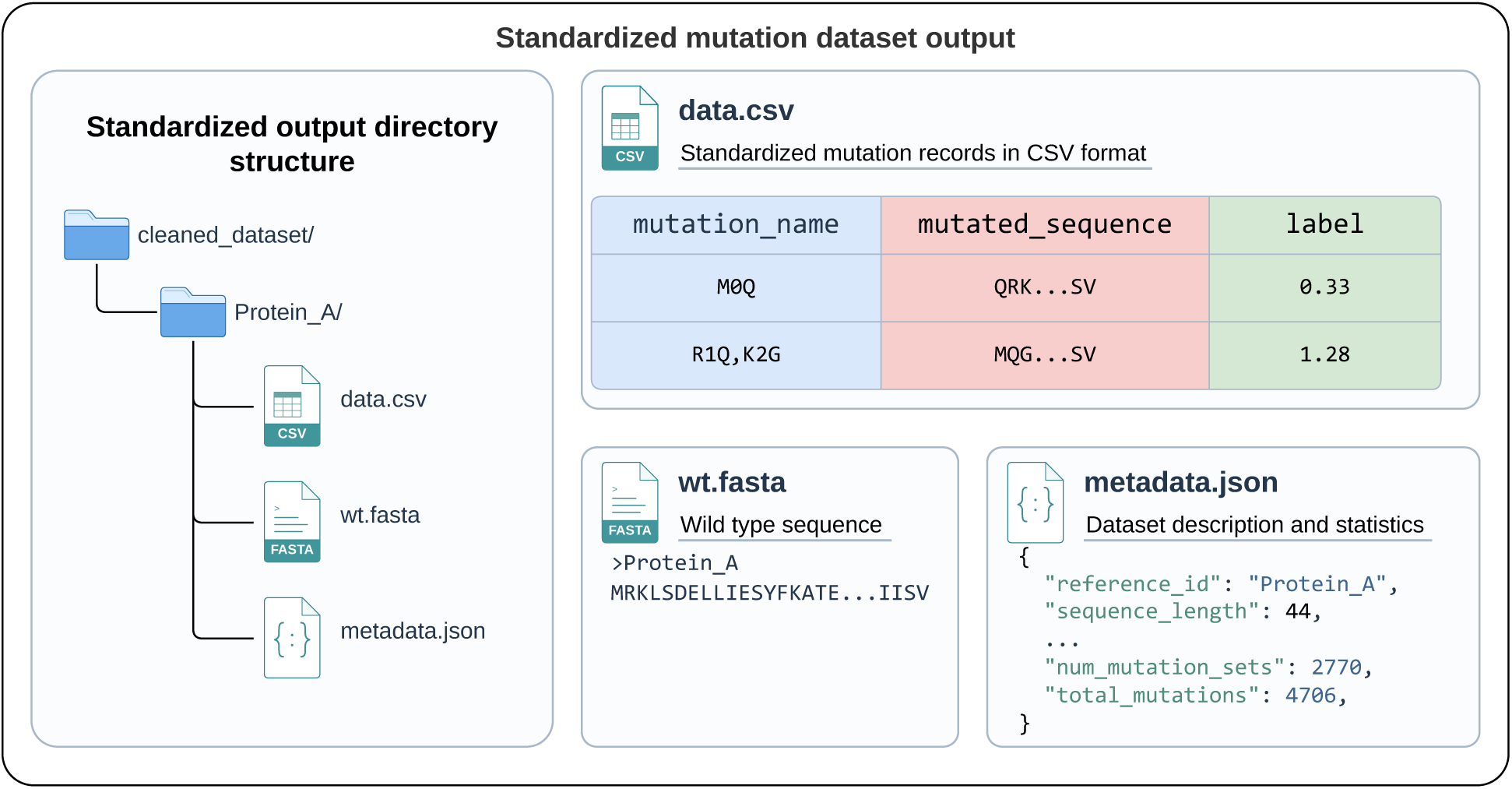
Standardized output structure generated by MutCleaner. Each cleaned dataset is organized by reference sequence and contains standardized mutation records (data.csv), the corresponding wild-type sequence (wt.fasta), and reference-sequence metadata and summary statistics (metadata.json).

### S3.1 data.csv

The data.csv file stores the standardized mutation records and corresponding label values associated with the reference sequence. It contains three columns: mutation_name, mutated_sequence, and label. The mutation_name field represents the standardized mutation annotation. A single amino acid mutation is represented in the form {wt_aa}{pos}{mut_aa}, where wt_aa and mut_aa denote the wild-type and mutant amino acids, respectively, and pos denotes the mutation position using 0-based indexing. For mutations involving multiple positions, individual mutations are ordered by position and separated by commas. The mutated_sequence field contains the mutated sequence obtained by applying the corresponding mutation set to the reference sequence. The label field stores the quantitative measurement associated with the mutation record.

### S3.2 wt.fasta

The wt.fasta file stores the reference sequence associated with the current subdirectory. For protein mutation datasets, the file contains the protein reference sequence; for codon-level DNA or RNA mutation datasets, it contains the corresponding DNA or RNA reference sequence. The FASTA header uses the sequence_name of the reference sequence when available; otherwise, the reference_id is used. Because each subdirectory corresponds to a single reference sequence, all mutation records in data.csv within the same directory are associated with the reference sequence stored in wt.fasta.

### S3.3 metadata.json

The metadata.json file stores basic information about the current reference sequence together with summary statistics for its associated mutation data. The reference_id field is the unique identifier used by MutCleaner for the reference sequence, whereas sequence_name stores its name. The sequence_type field specifies the sequence type and can be ProteinSequence, DNASequence, or RNASequence, and sequence_length records the length of the reference sequence. The position_unit field specifies the unit used for mutation positions. Protein mutations use residue as the position unit, whereas DNA or RNA codon mutations use codon. The num_mutation_sets and total_mutations fields record the number of mutation sets and the total number of individual mutations contained in those sets, respectively. The covered_positions and coverage_percentage fields indicate the number of positions affected by at least one mutation and the proportion of all reference positions that they represent, respectively, where the total number of positions is calculated according to position_unit. The num_unique_labels field records the number of distinct label values, has_unlabeled indicates whether any mutation records have missing labels, and dataset_name stores the name of the corresponding MutationDataset.

## S4 Curated dataset resources

MutCleaner cleaned and standardized 16 protein and codon mutation data resources comprising 12,143,621 mutation records in total. Summary statistics for each dataset are provided in Supplementary Table S1, including the numbers of reference entries, total mutation records, single-mutation records, and multiple-mutation records. Reference entries denote the number of independent standardized output directories in each dataset, with each directory corresponding to one reference sequence entry. Total, Single, and Multiple denote the numbers of all mutation records, single-mutation records, and multiple-mutation records, respectively.

**Table S1.** Summary statistics of raw and cleaned mutation datasets processed by MutCleaner.

| Dataset name | Sub-dataset name | Reference entries | Raw records | Total | Single | Multiple |
| --- | --- | --- | --- | --- | --- | --- |
| **dTm Dataset** | S557 | 39 | 571 | 557 | 557 | 0 |
|  | S4346 | 358 | 4,346 | 3,860 | 3,860 | 0 |
| **ddG Dataset** | S461 | 48 | 461 | 461 | 461 | 0 |
|  | S669 | 94 | 669 | 669 | 669 | 0 |
|  | S783 | 55 | 783 | 783 | 783 | 0 |
|  | S8754 | 305 | 8,754 | 6,411 | 6,411 | 0 |
|  | M1261 | 133 | 1,261 | 1,005 | 0 | 1,005 |
| **RBD ACE2 Dataset** | SARS-CoV-2-RBD Delta bc binding | 1 | 205,734 | 27,274 | 3,818 | 23,456 |
|  | SARS-CoV-2-RBD DMS variants bc binding | 3 | 385,031 | 39,314 | 11,283 | 28,031 |
|  | SARS-CoV-2-RBD DMS Omicron bc binding | 3 | 598,394 | 31,244 | 11,397 | 19,847 |
|  | SARS-CoV-2-RBD DMS Omicron-XBB-BQ bc binding | 2 | 412,404 | 13,607 | 7,633 | 5,974 |
|  | SARS-CoV-2-RBD DMS Omicron-EG5-FLip-BA286 bc binding | 3 | 512,560 | 23,913 | 11,438 | 12,475 |
| **MGnify ddG Dataset** | – | 7,794 | 65,710 | 65,710 | 8,408 | 57,302 |
| **TrpB Epistasis Dataset** | – | 1 | 194,041 | 159,999 | 76 | 159,923 |
| **RBD Antibody Dataset** | SARS-CoV-2-RBD MAP Moderna | 69 | 13,283,929 | 1,049,184 | 134,710 | 914,474 |
|  | SARS-CoV-2-RBD MAP Vir mAbs | 13 | 2,536,053 | 200,221 | 25,451 | 174,770 |
|  | SARS-CoV-2-RBD MAP Rockefeller | 53 | 10,339,293 | 814,470 | 103,738 | 710,732 |
| **CTXM Epistasis Dataset** | – | 2 | 292,410 | 79,848 | 646 | 79,202 |
| **Human Domainome Dataset** | Human Domainome Sup2 Dataset | 522 | 602,882 | 536,164 | 536,164 | 0 |
|  | Human Domainome Sup4 Dataset | 7,260 | 4,107,436 | 4,080,222 | 4,080,222 | 0 |
| **Codon cDNA Proteolysis Dataset** | – | 412 | 547,644 | 547,478 | 14,391 | 533,087 |
| **Antitoxin ParD3 Epistasis Dataset** | – | 1 | 9,261 | 7,999 | 57 | 7,942 |
| **Protein cDNA Proteolysis Dataset** | Protein cDNA Proteolysis dG Dataset | 479 | 776,298 | 582,865 | 426,965 | 155,900 |
|  | Protein cDNA Proteolysis ddG Dataset | 412 | 776,298 | 527,173 | 389,037 | 138,136 |
| **Codon DMS Substitutions Dataset** | – | 175 | 501,534 | 501,534 | 72,871 | 428,663 |
| **ArchStabMS1E10 Epistasis Dataset** | ArchStabMS1E10 Epistasis Sup4 Dataset | 5 | 428,172 | 356,934 | 120 | 356,814 |
|  | ArchStabMS1E10 Epistasis Sup5 Dataset | 5 | 3,202 | 3,033 | 153 | 2,880 |
| **ProteinGym DMS Substitutions Dataset** | – | 217 | 2,465,767 | 2,465,767 | 696,311 | 1,769,456 |
| **Codon Human Myoglobin Epistasis Dataset** | – | 1 | 9,714 | 9,112 | 4,073 | 5,039 |
| **Protein Human Myoglobin Epistasis Dataset** | – | 1 | 9,714 | 6,810 | 2,549 | 4,261 |

To quantify record-level quality-control outcomes during dataset curation, we counted five major categories of quality-control failures and record exclusions for each dataset: invalid or unsupported mutation annotations, invalid sequences, mutation–sequence inconsistencies, non-numeric labels, and missing values in fields required for standardized output (Supplementary Table S2). These categories are mutually exclusive, and each record was assigned according to the first corresponding check triggered during the cleaning workflow.

Mutation-related exclusions include both mutation annotations that cannot be correctly parsed or fail validity requirements and mutation types such as synonymous mutations, nonsense mutations, and addition or insertion mutations that fall outside the current scope of the standardized resources. Records in the latter group do not necessarily contain incorrect mutation annotations; rather, they were excluded because the currently curated resources are primarily intended for variant effect prediction models that use amino acid substitution mutations as input.

Sequence validity checks cover all input sequences required by the cleaning workflow for each dataset. The invalid-sequence category also includes cases in which sequence-coordinate information is incompatible with the corresponding sequence. For example, in the Human Domainome Sup4 Dataset, the PFAM_entry value ZN487_HUMAN/355-377 specifies an end position of 377, whereas the corresponding sequence has a length of only 207 residues. Records containing this type of out-of-range coordinate were therefore classified as invalid sequences.

A non-numeric label was defined as a value in the label field that was not encoded as NA but could not be converted to a numeric value, such as - or >6. Missing-value failures were restricted to fields required for standardized output that were explicitly encoded as NA. Mutation–sequence consistency was evaluated only after the sequence and mutation annotation had independently passed their respective validity checks.

**Table S2.** Dataset-wise counts of records excluded during MutCleaner processing.

| Dataset name | Invalid mutation annotations | Invalid sequences | Mutation–sequence inconsistencies | Non-numeric labels | Missing required values | Total validation failures |
| --- | --- | --- | --- | --- | --- | --- |
| TrpB Epistasis Dataset | 34,041 | 0 | 0 | 0 | 0 | 34,041 |
| RBD Antibody Dataset | 0 | 0 | 0 | 0 | 170,994 | 170,994 |
| Human Domainome Sup2 Dataset | 30,118 | 0 | 0 | 0 | 36,600 | 66,718 |
| Human Domainome Sup4 Dataset | 0 | 1,235 | 25,979 | 0 | 0 | 27,214 |
| Antitoxin ParD3 Epistasis Dataset | 1,261 | 0 | 0 | 0 | 0 | 1,261 |
| Protein Human Myoglobin Epistasis Dataset | 1,555 | 0 | 0 | 0 | 0 | 1,555 |
| Codon Human Myoglobin Epistasis Dataset | 602 | 0 | 0 | 0 | 0 | 602 |
| Protein cDNA Proteolysis dG Dataset | 86,416 | 0 | 0 | 100,831 | 0 | 187,247 |
| Protein cDNA Proteolysis ddG Dataset | 74,573 | 0 | 0 | 168,459 | 0 | 243,032 |

## S5 Parallel processing performance

To evaluate the runtime performance of MutCleaner under different parallel-processing configurations, we conducted the benchmark on a system equipped with an Intel Xeon Platinum 8352V 2.10 GHz CPU, 16 vCPUs, and 62 GB of memory, running Ubuntu 22.04.1 LTS and Python 3.13. The first 400,000 mutation records from the Protein cDNA Proteolysis Dataset were used as the test data, and data cleaning was performed using 1, 2, 4, 8, and 16 workers. Each worker configuration was run independently three times, and the mean runtime was used as the overall runtime for that configuration. Supplementary Fig S2 shows the runtime of MutCleaner with different numbers of workers. The points represent the mean runtime across the three independent runs, and the shaded region represents one standard deviation above and below the mean.

In addition to the mean runtime, we further used speedup to evaluate the performance improvement achieved by increasing the number of workers. For *p* workers, the speedup is defined as

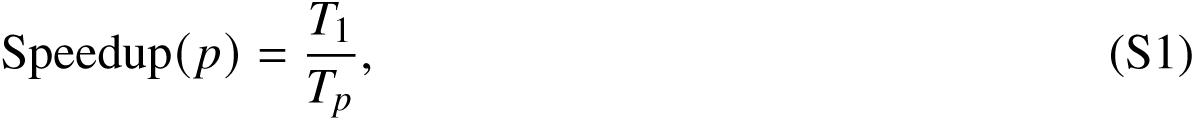

where *T*_1_ denotes the mean runtime using a single worker and *T_p_* denotes the mean runtime using *p* workers. A larger speedup indicates a greater improvement in runtime performance relative to the single-worker configuration.

The mean runtime and speedup for each worker configuration are shown in Supplementary Table S3. As the number of workers increased from 1 to 16, the mean runtime of MutCleaner decreased from 15.858 s to 12.410 s, corresponding to an increase in speedup from 1.000 to 1.278. These results show that increasing the number of workers reduced the data-cleaning runtime and improved the overall runtime performance of MutCleaner. However, the reduction in runtime became progressively smaller as the number of workers increased, indicating diminishing returns from additional parallelization.

**Figure S2.**
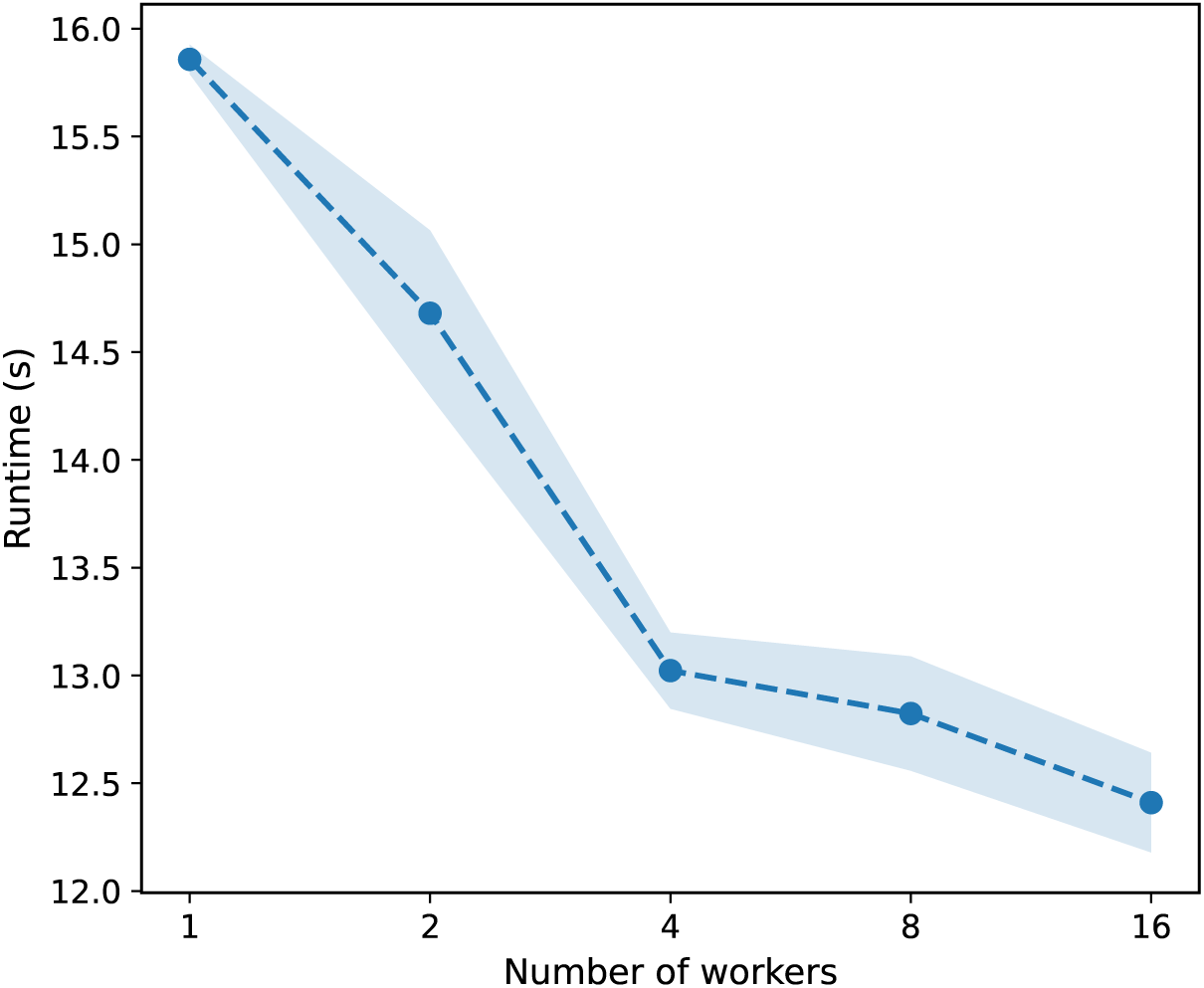
**Parallel processing performance of MutCleaner across different worker configurations.**

**Table S3.**
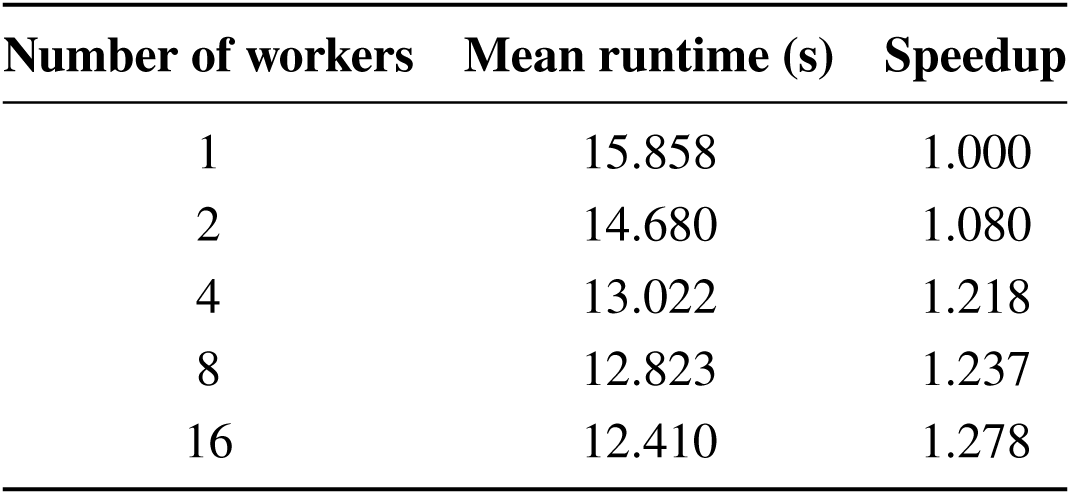
Mean runtime and speedup of MutCleaner with different numbers of workers.

## S6 Validation of MutCleaner data cleaning reliability

To evaluate the reliability and consistency of the MutCleaner data cleaning workflow, we compared the standardized datasets generated by MutCleaner with independently curated mutation datasets provided by ProteinGym. ProteinGym collects and standardizes protein mutation data from multiple published deep mutational scanning (DMS) experiments, and therefore serves as a reference dataset for evaluating the consistency of mutation data processing.

In this study, we selected DMS assays derived from the mega-scale cDNA-display proteolysis dataset established by Tsuboyama et al. [3], and compared the standardized results provided by ProteinGym with the results obtained after processing the original dataset using MutCleaner. For each ProteinGym assay, the corresponding MutCleaner output was identified based on the protein identifier contained in the assay filename. For example, the ProteinGym assay ODP2_GEOSE_Tsuboyama_2023_1W4G.csv corresponds to the 1W4G dataset in the MutCleaner output directory. The ProteinGym and MutCleaner datasets were matched using the standardized mutated_sequence field as the matching key. For each matched assay, the Spearman correlation coefficient and Pearson correlation coefficient were calculated between the ProteinGym DMS_score values and the MutCleaner label values.

In addition, to further evaluate the deviation between the experimental measurements from the two datasets, we calculated the Mean Absolute Label Difference (MALD):

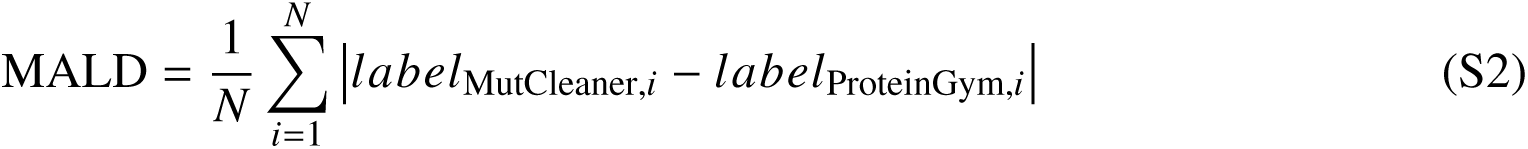

where *label*_MutCleaner_ represents the label values generated by MutCleaner, and *label*_ProteinGym_ represents the corresponding mutation effect values from ProteinGym (the DMS_score column). *N* represents the number of matched mutated sequences within each assay.

Overall, we obtained 64 matched DMS assays derived from the cDNA-display proteolysis dataset reported by Tsuboyama et al. and evaluated the data consistency between MutCleaner and ProteinGym for each assay. As summarized in Supplementary Table S4, the datasets processed by MutCleaner showed high agreement with the ProteinGym-curated datasets. Across all matched assays, the average Spearman and Pearson correlation coefficients were approximately 0.9996 and 0.9996, respectively, while the average Mean Absolute Label Difference was only 0.0032.

These results demonstrate that MutCleaner effectively preserves mutation information and experimental measurements during the data cleaning and standardization processes, and that its processed outputs are highly consistent with independently curated datasets, supporting the reliability of the MutCleaner data cleaning workflow.

**Table S4.** Validation of MutCleaner data cleaning consistency.

|  | **Spearman $\rho$** | **Pearson $r$** | **MALD** |
| --- | --- | --- | --- |
| **Average** | 0.9996 | 0.9996 | 0.0032 |

## S7 Roles and capabilities of ProteinGym, MaveDB, and MutCleaner in mutation dataset processing

Existing resources, including ProteinGym and MaveDB, have greatly facilitated the collection, standardization, and distribution of mutation effect datasets. However, these resources primarily focus on benchmark dataset construction or database management, whereas MutCleaner focuses on providing a reusable framework for cleaning, validating, and standardizing heterogeneous mutation datasets from diverse experimental sources.

To clarify the complementary roles of these resources, we compared the major functional capabilities of ProteinGym, MaveDB, and MutCleaner. ProteinGym primarily provides curated datasets for benchmarking variant effect prediction models, while MaveDB serves as a repository for storing and distributing multiplexed variant effect measurements. In contrast, MutCleaner provides a dataset-level preprocessing framework that converts heterogeneous mutation datasets into standardized resources with mutation and sequence consistency validation, failure tracking, and structured output formats.

Standardizing mutation annotations refers to converting heterogeneous mutation descriptions from different experimental sources into a consistent representation. ProteinGym performs dataset-specific processing during benchmark construction and provides standardized mutation annotations for released datasets. For example, multiple substitutions are represented using colon-separated mutation strings (e.g., A1G:B2V), and residue positions follow the conventional 1-based indexing scheme used in biological annotations. MaveDB defines standardized variant representations and validates submitted variants according to these rules. However, users generally need to preprocess heterogeneous mutation annotations before submission. In contrast, MutCleaner provides reusable functions for transforming diverse mutation annotations into a unified representation, including amino acid notation conversion, delimiter normalization, and indexing conversion. For example, multiple substitutions are represented using comma-separated mutation strings (e.g., A0G,B1V), with residue positions following 0-based indexing to facilitate downstream computational analysis.

**Table S5.** Comparison of functional capabilities among ProteinGym, MaveDB, and MutCleaner.

| **Capability** | **ProteinGym** | **MaveDB** | **MutCleaner** |
| --- | --- | --- | --- |
| **Primary purpose** | Benchmarking variant effect prediction models | Repository and distribution of MAVE datasets | Mutation dataset curation, cleaning, and standardization framework |
| **Curated mutation dataset resources** | ✓ | ✓ | ✓ |
| **Standardizing mutation annotations** | Dataset-specific standardization | Variant representation standard and validation | Built-in validation and standardization functions |
| **Validating sequence–mutation consistency** | Performed during dataset curation | Submission validation | Built-in validation functions |
| **Extensible preprocessing framework** | – | – | ✓ |
| **Tracking failed records** | – | – | ✓ |
| **Tracking cleaning step execution** | – | – | ✓ |
| **Providing inputs for variant effect prediction models** | ✓ | Requires additional preprocessing | ✓ |
| **Structured output format** | Mutation dataset CSV files | Database records with score tables and metadata | Mutation dataset CSV files, wild type sequence FASTA and metadata JSON |

Mutation–sequence consistency validation ensures that mutation annotations are compatible with the corresponding reference sequences. ProteinGym applies dataset-specific curation procedures during benchmark construction. We further examined ProteinGym datasets processed by MutCleaner and confirmed that the released mutation annotations were consistent with their corresponding mutated sequences. MaveDB performs submission-time validation of variants to ensure that deposited mutation representations are properly formatted and consistent with the target sequences. This validation is conducted during the data submission process to maintain the reliability of deposited datasets. In contrast, MutCleaner provides built-in validation functions that integrate mutation–sequence consistency checking into the preprocessing workflow. Given a wild-type sequence and mutation annotations, MutCleaner applies the mutations to the reference sequence to generate the corresponding mutated sequences. Successful generation of mutated sequences requires the mutation annotations to be consistent with the wild-type sequence; otherwise, the validation fails and the corresponding records can be identified for further inspection.

An extensible preprocessing framework enables users to adapt existing processing components and workflows to newly collected mutation datasets. MutCleaner provides an extensible preprocessing framework based on reusable cleaning functions and configurable pipelines.

Tracking failed records refers to the ability to preserve mutation records that do not pass validation or processing steps instead of directly discarding them. Retaining failed records enables subsequent data inspection, error analysis, and traceability during dataset cleaning. MutCleaner implements this capability by preserving records that fail during mutation parsing, sequence validation, mutation– sequence consistency checking, or other dataset-specific processing procedures. These failed records are exported as cleaning artifacts together with corresponding failure information, allowing users to identify problematic entries and analyze potential sources of data quality issues.

Tracking cleaning step execution refers to recording the status and runtime information of individual preprocessing steps during dataset cleaning. Unlike dataset-level metadata describing experimental information, execution tracking captures how a dataset is processed. MutCleaner records the execution status of each cleaning step, including step names, completion status, runtime information, and error messages. These records enable users to inspect the cleaning process, identify failures, and improve reproducibility of dataset preprocessing.

Sequence-level inputs for variant effect prediction models typically include standardized mutation annotations, corresponding mutated sequences, and experimentally measured mutation effects that can be directly used for model training or evaluation. MaveDB provides deposited variant effect measurements together with variant representations and associated metadata. However, the deposited variants usually require additional preprocessing by users, such as converting mutation annotations into mutated sequences before being used in sequence-based prediction models.

Structured output format refers to the organization of standardized mutation datasets and associated information for downstream analysis. ProteinGym primarily distributes curated benchmark datasets as standardized mutation tables, where each dataset is provided as a CSV file containing mutation annotations and corresponding experimental measurements. MaveDB organizes mutation effect datasets as database records. Each deposited score set contains variant information, score tables, target sequence information, and associated metadata, enabling standardized storage and distribution of MAVE measurements. In contrast, MutCleaner exports each processed reference sequence as an independent standardized dataset package containing three major files: data.csv, wt.fasta, and metadata.json. The data.csv file stores standardized mutation annotations, mutated sequences, and experimental labels; the wt.fasta file provides the corresponding reference (wild-type) sequence; and the metadata.json file records reference-sequence metadata and summary statistics. This sequence-aware output organization facilitates direct reuse of cleaned datasets in downstream computational analyses.

